# *ORIGAMI*: A Software Suite for Activated Ion Mobility Mass Spectrometry (aIM-MS) Applied To Multimeric Protein Assemblies

**DOI:** 10.1101/152686

**Authors:** Lukasz G. Migas, Aidan P. France, Bruno Bellina, Perdita E. Barran

**Author notes:** Correspondence and request for materials should be addressed to Professor Perdita Barran.

## Abstract

We present here a software suite (**ORIGAMI**) that facilitates the rapid acquisition and analysis of ion mobility data following collisional activation. **ORIGAMI** was developed for use on Waters Synapt instruments where data acquisition is achieved by interfacing WREnS (Waters Research Enabled Software) and MassLynx. Two components are presented, the first is **ORIGAMI***^MS^* which enables activation of ions by sequential increase of collision voltages prior to ion mobility analysis. We demonstrate the use of **ORIGAMI** on the tetrameric assemblies formed by the proteins concanavalin A (103 kDa) and alcohol dehydrogenase (143 kDa). Activation is performed in the trap collision cell of the Synapt TriWave assembly, where the collision voltage can be ramped from 0-200 V. All of the acquired data is recorded in a single file which simplifies data acquisition. This substantially decreases the time needed to perform a typical activated IM-MS experiment on a single protein charge state from approx. 2 hours to ~25 minutes. Following data acquisition the data is analysed in the second component, **ORIGAMI***^ANALYSE^*, which allows the user to visualise the effect of activation on the mobility of the parent ion, as well as on any produced fragment ion. The user can export the data in the form of heat maps, waterfall or wire plots. In addition, tools implemented in **ORIGAMI** enable easy data extraction from single or multiple MassLynx .raw files, in-depth interrogation of large datasets, statistical analysis and figure creation capabilities. We demonstrate the use of **ORIGAMI** on concanavalin A and alcohol dehydrogenase acquired using the traditional protocols and the **ORIGAMI*^MS^*** method.

**Highlights:**

- New methodology for faster acquisition and data analysis following activation of ions separated by ion mobility mass spectrometry
- Software package capable of simultaneous analysis of multiple MassLynx .raw files
- Visualization of the change in mobility of parent and fragment ions following activation
- Easy extraction, data processing and extensive plotting tools

## Introduction

Mass spectrometry (MS) and ion mobility mass spectrometry (IM-MS) are commonly used to characterize small molecules,^1,2^ peptides,^3^ proteins^4^ and their complexes.^5^ IM-MS has an increasing relevance to structural biology, and whilst MS alone can provide information about the molecular weight, charge state distribution and stoichiometric information of the sample, the addition of IM separation into a workflow can help elucidate the size, shape and structural stability of a given analyte. IM-MS has been utilised to study the conformational landscapes of mono- and homo-oligomeric proteins,^6^ protein-protein^7^ and protein-ligand complexes.^4,8–10^ In addition to gas phase separation, MS and IM-MS has been used to study biophysical phenomena such as gas phase protein rearrangement,^8^ stabilisation and rearrangement of protein ensembles upon ligand binding,^8,11^ proton- and electron transfer mechanisms^12,13^ and protein unfolding^4,14,15^, with the latter becoming increasingly more popular.

In IM-MS, conformations are determined by measuring the time it takes ions to cross a mobility device and recording an arrival time distribution (ATD) which can then be converted to a rotationally averaged collision cross section (CCS). The separation is enabled by pulsing packets of ions into a mobility cell filled with an inert buffer gas held under the influence of a weak electric field, the latter serving to propel the ions through the cell with a velocity that depends on their charge and on the nature of the collisions with the buffer gas.^16,17^ Compact ions undergo fewer collisions with the gas and will exit the cell earlier than more extended forms. Presently, the majority of IM-MS research on biomolecular complexes in academia and in industry is carried out on Waters Synapt^17,18^ instruments (G1, G2 and newer) fitted with travelling wave IM-MS (TWIMS-MS) which can record the ATD for each ion. However, to obtain rotationally averaged collision cross sections (CCS), the TWIMS measurements need to be calibrated against samples with known CCS.^19^ Conversion of ATD to CCS is not strictly necessary, but it does remove the influence of ion charge, drift gas pressure and temperature, and other experimental variables on the data and is therefore certainly beneficial for inter-lab comparison. CCS can be ‘learnt’ for any static structure using *ab initio* methods to mimic the experiment or data scaled from empirical values, and/or machine learning; here also having experimental data in the form of CCS values facilitates comparison. For experiments that are using IM-MS to examine changes in conformation of closely related molecular ions, for example *apo* and *holo* forms of proteins or mutated *vs*. wild type, simply comparing IM-MS data taken on the same instrument can be highly informative.^4,20,21^

The process of gas phase protein unfolding monitored by mobility measurements, which recently has been referred to as Collision Induced Unfolding (CIU), but perhaps more accurately can be termed activated ion mobility mass spectrometry **a**IM-MS. An early demonstration of aIM-MS on proteins was conducted by Shelimov *et al*.^22^ who investigated the stabilising influence of disulfide bonds by comparing the behaviour of BPTI (three disulfide bonds) with cytochrome *c* (none). Following these early experiments, aIM-MS was used in some analytical workflows but overall output remained low,^23,24^ in line with the lack of commercial IM-MS instrumentation. Hopper *et al*. used a trap ion guide to kinetically energise protein-ligand complexes and be able to differentiate species based on their ATDs at elevated trap voltages.^14^ More recently, aIM-MS methodology has been applied to more complicated systems to study the unfolding mechanisms of mono-, di- and tetrameric proteins,^25,26^ differentiate biosimilars^27,28^ and observe alterations to protein stability upon single point mutation^21^ and ligand and cofactor binding.^4,20,29^

**a**IM-MS in a Synapt mass spectrometer is facilitated by sequentially increasing the potential difference between the source and trap region of the instrument which effectively raises the ions kinetic energy in the presence of the trapping gas, followed by IM-MS separation which can then be used to monitor the influence of activation on the ion of interest. Unsurprisingly, as an ions kinetic energy increases, its conformation can adapt in order to accommodate its excessive internal energy prior to fragmentation. This process often leads to a marked change in the recorded ATD. Despite IM-MS being a relatively low resolution technique in comparison to other biophysical methods (*i.e*. X-ray crystallography or NMR), it offers multiple benefits including low sample consumption, relatively quick acquisition and analysis times and the potential for high throughput screening. In spite of these benefits, when applied to large macromolecular complexes, such as proteins, aIM-MS methodology contains several bottlenecks that preclude its wide adoption. The current barriers include: (i) sequential ramping of collision voltages (CV) is monotonous, time-consuming and prone to errors since the MS operator needs to manually alter the CV; (ii) current protocols result in multiple files and large volumes of data handling and (iii) extraction of ATDs for an ion at given CVs is time consuming, not systematic and easily varies from user to user. While there are existing open-source software packages capable of data visualisation (*CIUSuite*)^30^ and efficient data extraction (*PULSAR*),^31^ these programs are not designed for, or are capable of offering efficient aIM acquisition or high-throughput analysis. In this report, we present a novel open-source software package for semi-autonomous CIU data acquisition (**ORIGAMI***^MS^*) for Synapt G2, G2S and G2S*i* instruments, with our complementary rapid data analysis tool (**ORIGAMI***^ANALYSE^*). The acquisition software automates the aIM process by sequentially raising the collision voltage within a single MassLynx file, whilst **ORIGAMI***^ANALYSE^* enables its direct analysis, reducing the acquisition time scales, quantity of data handling and analysis time. The software is also back compatible to enable the analysis of manually acquired datasets and text files. The primary role of **ORIGAMI** is to automate the acquisition and thus reduce analysis time. **ORIGAMI***^MS^* reduces the experimental footprint (number and size of files) and improves the reproducibility and reliability of aIM-MS protocols. In order to demonstrate the capabilities and benefits of using **ORIGAMI**, we present its use on two multimeric proteins that are readily available as standard molecules.

## Materials and methods

### Materials

Concanavalin A (ConA, jack bean), alcohol dehydrogenase (AdH, *Saccharomyces cerevisiae*), and ammonium acetate were purchased from Sigma (St. Louis, MO, USA). All protein samples were buffer exchanged into ammonium acetate at pH 7.4 using Micro Bio-Spin 6 columns (Bio-Rad, Hercules, CA, USA) and prepared to a final concentration of 10 μM (ConA, AdH; 200mM AmAc). Either fresh or flash frozen samples thawed on ice were used for analysis

### Experimental conditions

All experiments were carried out on an unmodified Synapt G2 and G2*S* (Waters, Wilmslow, UK) in positive ionisation mode with nitrogen as the buffer gas. Samples were ionized by applying a positive potential through a platinum wire (Diameter 0.125 mm, Goodfellow) inserted into nESI tips that were pulled in-house from thin-walled glass capillaries (i.d. 0.69 mm, o.d. 1.2 mm, Sutter Instrument Company, Novato, CA, USA.) using a P2000/F laser puller (Sutter Instrument Co., Novato, CA, USA). All ions were generated via electrospray ionisation using a capillary voltage of 0.7-1.2 kV, a sampling cone of 10-20 V and a source temperature of 40 °C. All other experimental voltages (including travelling wave settings) were minimised in order to reduce protein activation prior to the activation process. Detailed parameters are shown in SI Table 1. In order to raise the transmission efficiency of ConA and AdH, the backing pressure was set to 5-7.5x10^1^ mbar and the Trap cell gas flow to 4-7 mL/min.

Collisional activation of analytes was accomplished by sequentially increasing the collision voltage in the Trap collision cell prior to ion mobility separation using both **ORIGAMI***^MS^* and the manual protocols. The collision voltage was typically ramped from 4 to 200 V in 2 V increments. Detailed parameters of the aIM-MS process are shown in SI Table 2. An exemplar charge state ([AdH+24H]^24+^ and [ConA+20H]^20+^ for each protein was first mass selected in the quadrupole prior to activated ion mobility separation; mass spectra and arrival time distributions were collected in MSMS mode.

### Software design

**ORIGAMI** is an open-source software package consisting of two components, **ORIGAMI***^MS^*, capable of streamlined aIM-MS data acquisition and **ORIGAMI***^ANALYSE^*, used to extract, analyse and plot activation fingerprint data. The software was developed using Python 2.7 programming language utilising several popular modules, namely *Numpy*,^32^ *SciPy*, *Matplotlib*^33^ and *Mayavi*^34^ amongst many others. A graphical user interface (GUI) was developed in wxPython. Both programs are distributed as pre-compiled executables available from https://www.click2go.umip.com/i/s_w/ORIGAMI.html free of charge for academic use.

The autonomous **ORIGAMI***^MS^* software was developed in *C#* using the Waters Research Enabled Software (WREnS), which enables additional controls of the Synapt hardware. In order to achieve controlled collisional activation independently of MassLynx, WREnS is here used to control ion transmission by modulation of DC potentials, in particular the DRE lens to control the ion signal attenuation and ion source and trap collision cell voltages. While no hardware modifications are required to utilise **ORIGAMI***^MS^*, the user is required to install WREnS on their instrument PCs. WREnS can be obtained by research agreement from Waters (Wilmslow, UK) with installation instructions. **ORIGAMI***^MS^* can be used in three ways, utilising the user-friendly GUI (shown in Figure S1), executed from the command line or as a pre-process program in the MassLynx worklist.

The analysis software, **ORIGAMI***^ANALYSE^* accepts several data types, including MassLynx raw files, text files and correctly formatted Microsoft Excel spreadsheets. Rather than ‘reinventing the wheel’ we have adopted components of the Driftscope software (Waters, Wilmslow, UK) to read MassLynx .raw files, supplied with the Synapt hardware. Driftscope enables quick extraction of the IM-MS data in a binary format which can be readily converted to a [*n* x 200] matrix, where *n* refer to the number of scans, which in turn may track a collision voltage ramp or indeed any other activation or ion mobility separation parameter. The program has no specific pre-requisites, however in order to extract MS and IM-MS data from MassLynx .raw files, a local installation of Driftscope is required. Presently data extraction is only available on Windows platforms as for all other analyses of MassLynx data. The analysis software was built and tested on Microsoft Windows 7/8/10 with 64-bit hardware architecture, however Linux and Apple Mac OS X versions are in development.

**ORIGAMI***^ANALYSE^* can import either single or multiple MassLynx files simultaneously, providing the same analysis support for both manually and autonomously acquired datasets. Regardless of the input format, mass spectra are loaded as a sum of all scans for each loaded MassLynx file. In the case of *ORIGAMI^MS^* acquired data, the default displayed mass spectrum is a composite of all scans within the file consisting of all collision voltage increments; individual mass spectra for specific collision voltages can be extracted and saved separately in a text format. Initially, the 2D/3D aIM plot is shown for the entire mass spectrum, typically consisting of a combination of precursor and fragment ions. In order to examine single charge states, the user can select a *m/z* region of interest permitting the corresponding ATD profile for each collision voltage to be extracted and stitched together to form a aIM matrix. **ORIGAMI***^ANALYSE^* is capable of mining multiple ions simultaneously and saving them in a text or compressed binary format for storage or further analysis in other software packages. Figure S2 shows the graphical user interface used to analyse **ORIGAMI***^MS^* acquired dataset, utilising the multi-ion selection tool. Figure S3 shows similar user interface, used to analyse a list of text files.

## Results

### The ORIGAMI*^MS^* workflow

The backbone of the **ORIGAMI** package is **ORIGAMI***^MS^*, which has been designed as a user-friendly program to develop advanced mass spectrometry workflows. Here it is used to automate an increase in applied collision voltages to induce ion restructuring and/or dissociation. On Waters Synapt instruments, collisional activation can occur prior to mass selection in the sampling cone, post the quadrupole in the trap collision cell, or post TWIM separation in the transfer collision cell, on either mass selected or all molecular ions. Here we only explore activation in the trap collision cell, but we have also implemented a mode for sampling cone activation, which is more suited to analysis where the analyte presents in a single charge state.

The collision voltage is slowly raised from CESTART to CEEND with a small increment CESTEP. In **ORIGAMI***^MS^*, the user can specify the region of the mass spectrometer where collisional activation is to be performed (*i.e*. cone or trap), ion polarity, starting and ending voltage, step size, scan time and time spent on each collision voltage. Once an acquisition is started, the collision voltage ramp is performed autonomously and the data is recorded to a single MassLynx file.

A simplified example of the **ORIGAMI***^MS^* workflow is shown in Figure 1a. In this exemplar case, the area shown in red depicts the collision voltage profile as the voltage is raised. Here, the collision voltage is raised from 5 to 35 V with an increment of 5 V. Each collision voltage is kept constant for 3 scans with a scan time of 5 seconds, resulting in 15 seconds of acquisition per voltage (Figure 1a inset). The two areas shown in blue (before and after the CE ramp) represent ‘reporter’ periods in which the ion flow is stopped in order for **ORIGAM**I*^ANALYSE^* to determine when the activation process has started and stopped. The areas shown in green represent the initial experimental conditions which are reset at the end of the aIM-MS protocol. The procedure shown in Figure 1a gives a total data acquisition time of 2.5 minutes, however in most experiments the voltage ramp will span a wider range of values. The approximate acquisition times shown in Figure 1b are for experiments using voltage increments of 2 V with 3 scans per voltage and a scan time of 5 seconds. Typical experimental parameters and their ranges are shown in Figure 1c.

**Figure 1.**
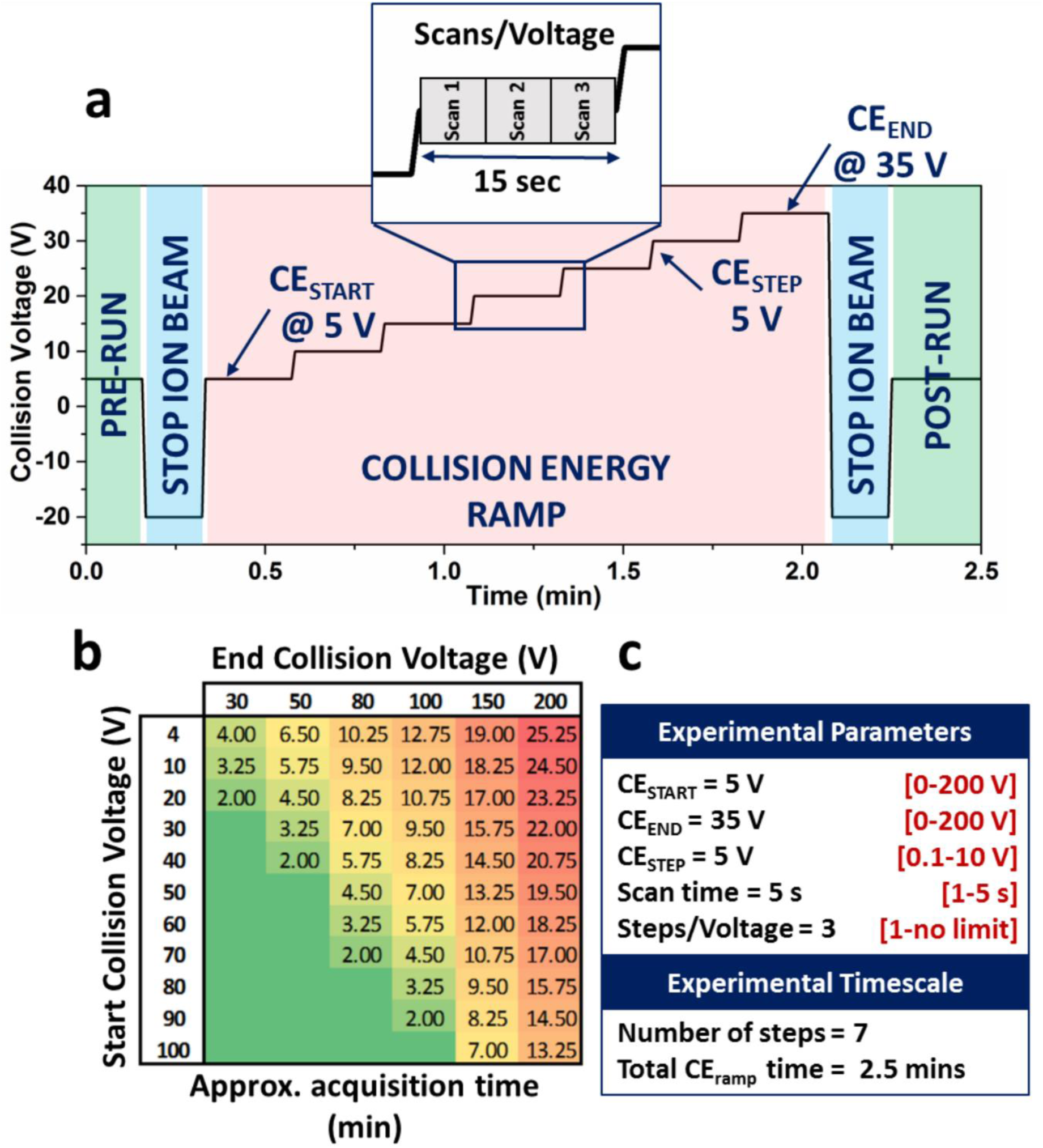
Procedure for activated IM-MS with a Collision Voltage ramp carried out using **ORIGAMI***^MS^*. (a) Schematic to show the sequence of a typical experiment wherein the collision voltage is raised from 5 V to 35 V with a 5 V increment. In this example, data is acquired for three scans at each collision voltage with a pre-set scan time of 5 s (15 s per 5 V, shown in inset). The area highlighted in green represents the initial experimental conditions (before and after the aIM-MS experiment) and the blue area shows the two periods for which the ion signal is attenuated to < 1 % of its original value in order to delineate when the aIM-MS process has started and finished. The entire experiment in this example takes ~2.5 min. (b) a reference table which shows the approximate acquisition times based on a pre-set scan time of 5 s, with a CE increment of 2 V and 3 scans per voltage. (c) Typical experimental parameters and the minimum/maximum ranges.

Often as a consequence of increasing the collision voltage, the signal intensity of the precursor ion decays rapidly. In order to account for the decrease in the ion intensity we have developed three methods where the number of scans per voltage (SPV) increase as a function of the collision voltage. The original method is referred to as *linear* (shown above), whilst the more advanced methods include *exponential, fitted* and *user-specified;* these methods have been developed *via* extensive aIM-MS studies encompassing a range of biomolecules (data not shown). Briefly, the *exponential* method increases the number of SPV as a product of the exponential function scaled by two user-specified parameters which define the relative starting point and rate of the SPV rise. The *fitted* method is based on a modified Boltzmann equation fitted to multiple datasets that represent the common ion intensity decay rate amongst multiple biomolecules. The *fitted* method accepts one user-specified parameter which defines the rate of SPV increase. Finally, the *user-specified* method accepts a list of SPVs for each collision voltage supplied by the user. These non-linear ramp functions can improve the signal-to-noise for the precursor and fragment ions but also lead to longer acquisition times.

### ORIGAMI*^ANALYSE^* overview

**ORIGAMI***^ANALYSE^* was developed to tackle one of the major bottlenecks of the activated IM-MS protocol, namely the time-consuming data processing, as well as to enhance data visualisation. Our aims were: to create software capable of reading MassLynx and text files; to accelerate data processing; and to allow the user to better view the acquired data which in turn will assist interpretation of the experiment performed. A summary of functions found in the program is shown in Figure 2 and discussed in more detail below. The majority of built-in methods can operate on MassLynx, text and Excel files, with the obvious exception of *m/z* data extraction. A typical workflow associated with using **ORIGAMI***^MS^* and **ORIGAMI***^ANALYSE^* is shown in Figure S4, in this case for alcohol dehydrogenase 24+. The software can be operated in two ways, using the GUI or in a high-throughput batch mode using the command line. 2D and 3D visualisation of **a**IM datasets are the most informative way of looking at activated analyte fingerprints. These plots convey information about the ATD profiles at individual collision voltages, adequately display the relative intensity of conformational families and easily highlight trends and unfolding motifs. The visualisation window offers easy navigation and a high level of interaction, enabling detailed analysis of specific regions of the **a**IM fingerprints. The raw data can be normalised, smoothed, interpolated and filtered within **ORIGAMI***^ANALYSE^* in order to improve the image quality.

**Figure 2.**
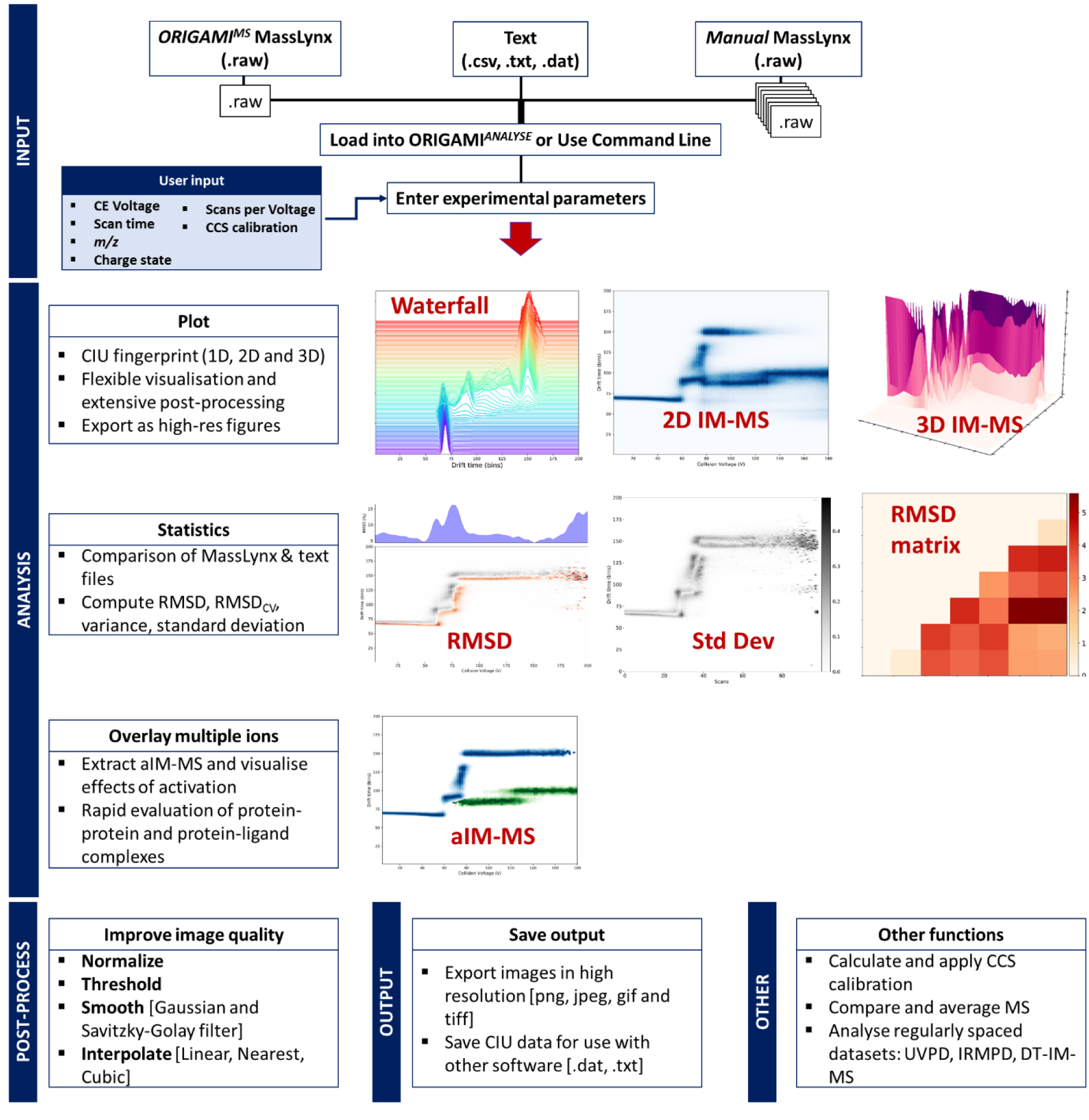
General schematic of features available in **ORIGAMI***^ANALYSE^*.

### aIM Fingerprint comparison

A common visualisation technique to observe structural, conformational and stability differences between samples is to use a subtraction heatmap and root mean square deviation (RMSD) calculation. These can be used to quantify the difference between datasets by subtracting two aIM matrices and computing the pairwise difference. The RMSD can be calculated using equation 1:

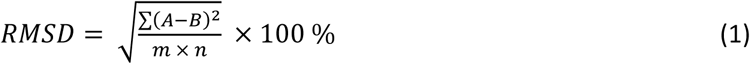

where A and B are aIM data matrices of identical size with the dimensions of [*n* x *m*] (*n* is the number collision voltage points and *m* is the number of IM-MS bins). Another way to visualise the RMSD contour map is to plot the RMSD as a function of the collision voltage (RMSD_CV_), calculated using equation 2:

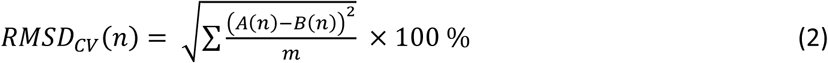

where A and B are aIM lists for individual *n* with the dimensions of *m*. RMSD plots (Figure 3b) are more sensitive to ‘global’ differences in the collisional unfolding of analytes and can easily discriminate between permutants, single and multi-point mutations or the effects of ligand binding on protein stability. In contrast, the RMSD_CV_ (Figure 3a) offers a ‘local’ insight into the motifs as it examines the minor fluctuations occurring at each collision voltage. The RMSD_CV_ can help identify differences at specific collision voltages and is most suited for day-to-day and batch-to-batch comparisons in addition to applications mentioned above.

**Figure 3.**
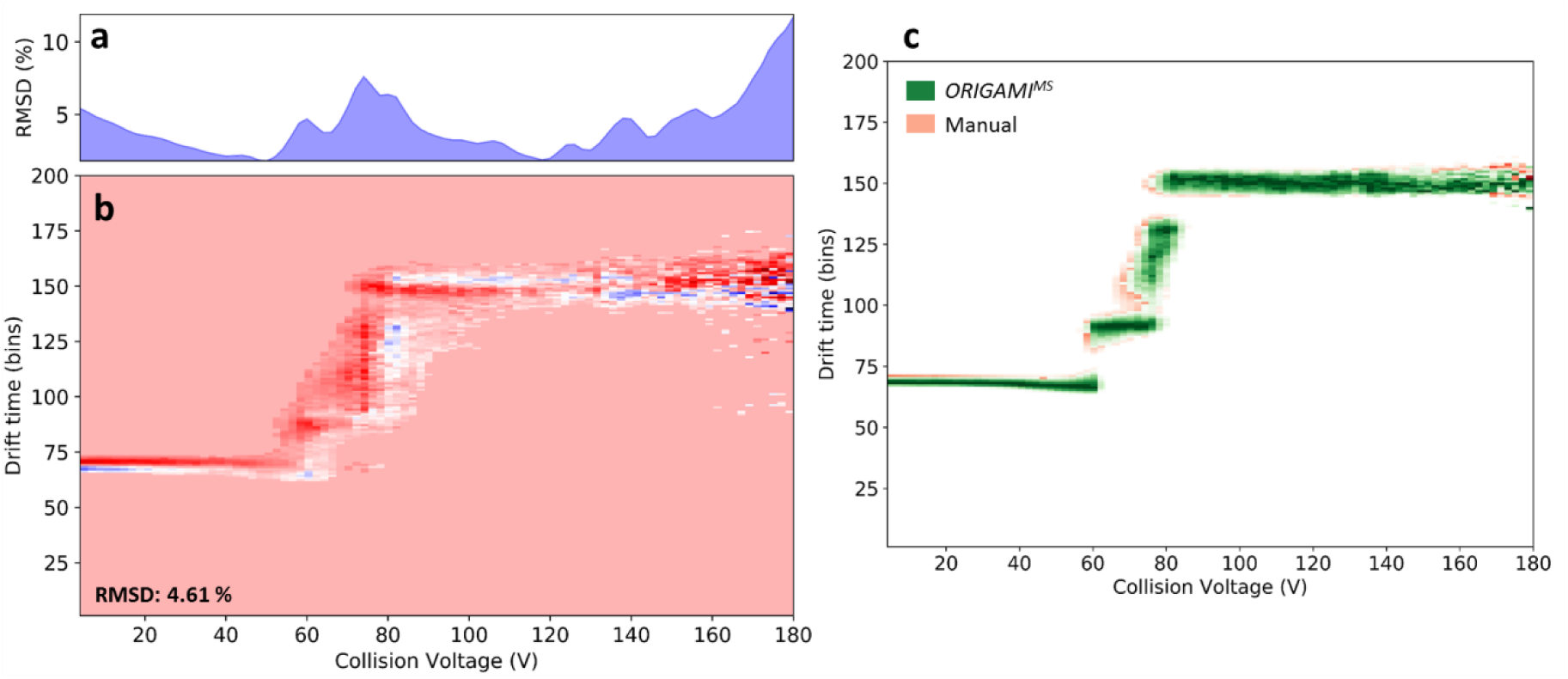
Comparison of manually and **ORIGAMI***^MS^* acquired datasets spanning identical collision voltage ranges for the ConA [4M+20H]^20+^ ion. (a) RMSD_CV_ plot highlighting collision voltages with most significant differences in their arrival time distributions and relative intensities. (b) A subtraction plot of two matrices with an overall RMSD error of 4.61 %. (c) An overlay plot of the two datasets with manually acquired data shown in red and **ORIGAMI***^MS^* in green. The plots were generated by applying a mask of different colour to each dataset individually.

While the usage of RMSD values and subtraction plots is unequivocally useful, the technique is reliant on the two datasets being of identical size and encompassing the same collision voltage range. In the case where *n* or the individual collision voltages differ, the subtraction plot will be misaligned and the corresponding RMSD values greatly exaggerated; minor shifts in the ATD profile which can be caused by a number of external factors (*i.e*. differences in IMS or trap pressures, source conditions, or intermittent spray) can lead to over-stated RMSD differences. In order to alleviate some of these issues, the aIM data matrices are typically averaged over three replicates obtained on separate days, allowing the average aIM data fingerprints to be compared. In most cases the ion of interest occupies a fraction of the aIM data, hence the matrix can be cropped or a mask (threshold) applied to remove noise peaks, this however tends to increase the RMSD values. Typical image processing methods, such as image smoothing, noise suppression and image interpolation can be used and have significant impact on the RMSD score and the subtraction plot.

Another method to visualize the aIM fingerprints is to overlay two (or more) matrices and represent them as individually coloured masks (shown in Figure 3c). While the overlay method provides no statistical or numerical information about the difference (or similarity) of the two analytes, it enables direct comparison of multiple matrices and offers clear recognition of regions of interest (also shown in Figure 4c).

**Figure 4.**
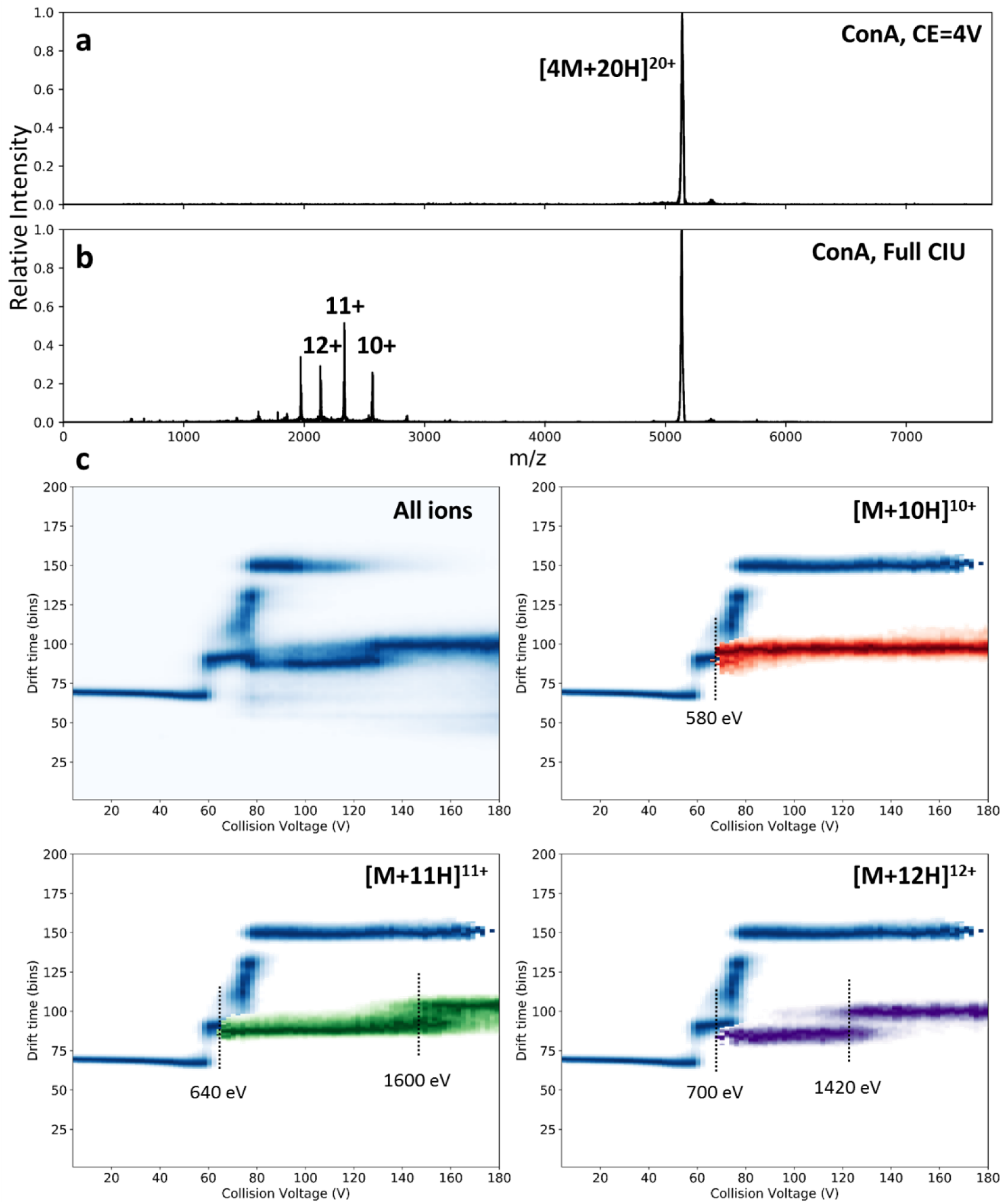
Output from **ORIGAMI***^MS^* acquisition following **ORIGAMI***^ANALYSE^* mediated aIM-MS on the tetrameric protein Concanavalin A [4M+20H]^20+^. The data was acquired using a linear voltage ramp from 4-180 V with a step size of 2 V and 15 s of acquisition per collision voltage (3 SPV and scan time of 5 s) and a total experiment time of 23 minutes. (a) Mass spectrum taken at 4 V showing the presence of only ConA [4M+20H]^20+^; (b) MS taken from the entire aIM-MS acquisition showing the presence of precursor and fragment ions (primarily monomers); (c) aIM-MS data of all ions and three of the product ions assigned to the [M+10H]^10+^ (red), [M+11H]^11+^ (green) and [M+12H]^12+^ (purple) monomers of ConA overlayed on the aIM-MS data for the [4M+20H]^20+^ parent ion (blue). The monomeric ions appear at 58 V following first major unfolding event.

### aIM fragment overlay

Information obtained from a single activated IM-MS experiment can complement non activated IM-MS and non IM tandem mass spectrometry experiments by providing detailed information about ion unfolding pathways, stabilities and conformational diversity; with the implementation of **ORIGAMI***^MS^* into the analytical workflow, the collision voltage ramp can be fully controlled and accomplished with speed and ease. Unsurprisingly during collisional activation, the kinetically excited precursor ion can undergo CID fragmentation, which is dependent on multiple parameters including inter- and intramolecular bonds, applied collision voltage and ion charge state.^26^ For large macromolecular complexes, collisional activation can cause unfolding, and fragmentation patterns may be a readout of structural rearrangements. Whilst it is common in tandem mass spectrometry to present the fragment ions, to date aIM-MS work has principally considered the effects of activation on the precursor ion, with an experimental workflow that normalises the data from each collision voltage. Since this later approach does not present all of the information from the experiment, for instance the unfolding pathways of fragment ions, we have developed **ORIGAMI***^ANALYSE^ to* address this. **ORIGAMI***^ANALYSE^* allows direct, facile extraction of multiple ions from MassLynx raw files (manual and **ORIGAMI***^MS^*), the subsequent visualisation of the precursor and fragment ions in 2D/3D enables a more appropriate interrogation of the relationship between molecular structure and ion dissociation pathways. In addition, by integrating TWIMS calibration into the workflow, the collision cross section data can be directly used to quantify the structural changes occurring during the unfolding and dissociation processes. Moreover, integration of the methodology employed by Allison *et al*.^31^ whereby, protein and protein-ligand stabilities were extracted from gas phase aIM-MS experiments, could provide useful insights which can be utilised to aid conformational sampling techniques.

To demonstrate the aIM-MS overlay method for biomolecules, we show the collisional unfolding of the ConA 20+ tetramer obtained using **ORIGAMI***^MS^*. Activated IM-MS was performed between 4-180 V with a step size of 2 V and 3 scans per voltage (15 s). The ConA ion retains a narrow ATD profile and collapses slightly until the collision voltage increases to 58 V (1160 eV) where the first unfolding event occurs.

Upon disruption of the non-covalent interactions, the first monomeric ions appear. Figure 4a shows the mass spectrum at collision voltage of 4 V, showing only the precursor ion peak; Figure 4b shows the total mass spectrum obtained for the entire aIM-MS experiment, where the monomeric fragment ions produced by activation are observed in a charge state range of 9+ to 14+ as has been reported previously for non IM tandem MS. Since unfolding and fragmentation occurs in the trap prior to mobility separation, fragment monomeric ions can undergo further unfolding and even MS^3^. A 2D aIM heatmap for all ions is shown in Figure 4c (top left), whilst an overlay of the 3 dominant monomer product ions along with the tetrameric parent species is shown alongside As previously reported, collisional unfolding pathways are dependent on the charge state of the ion;^26^ in the case of the 10+ ion (top right, red), the species appear upon the first unfolding event and the protein retains the ‘original’ unfolded state without further structural rearrangement. The 11+ (bottom left, green) and 12+ (bottom right, purple) ions are also released at the same time, however these experience further unfolding at 146 V (1600 eV) and 118 V (1420 eV), respectively. The utilisation of the overlay method and rapid extraction tools can significantly enhance the analytical workflows to monitor precursor and fragment ions. An immediate candidate for such protocol is to monitor non-covalent binding interactions of cofactors or drug candidates to biomolecules.

### TWIMS calibration

Determining collision cross sections from TWIMS measurements requires calibration using appropriate ions with known CCS values;^35,36^ the calibration procedure is necessary due to the non-linear relationship between the mobility and drift time of the ions separated by the travelling wave. Since the first release of Waters Synapt instruments, several calibration protocols have been developed, all of which rely upon the use of one or more calibrant ions selected based on the analyte type and molecular weight.^19,37,38^ As with many analytical measurements, their accuracy is determined by the selection of appropriate calibrants and careful measurements. Consequently, the experimental conditions (particularly regarding IM and ToF parameters) for both the unknown and calibrant ions must be kept constant.

**ORIGAMI***^ANALYSE^* automates the calibration procedure by loading a MassLynx file with calibrant data, extracting the mass spectrum, assigning the charge states to the present peaks and extracting their arrival time distributions. The program automatically detects the apex of the ATD and matches it with the appropriate CCS value. The resultant calibration curve can be modified by simply selecting or deselecting desired ions. Creating calibration curves with more than one calibrant is also possible by loading multiple MassLynx files. Alternatively calibration curve parameters obtained from elsewhere (*i.e*. manually calibrated) can also be used. The resultant calibration parameters can be used on the **ORIGAMI***^MS^* or manually acquired datasets.

## Conclusions

**ORIGAMI** provides an efficient means of data acquisition and more in-depth analysis of TWIMS acquired aIM datasets. The automation of the collision voltage ramp can be utilised to use aIM in high-throughput environments, effectively reducing acquisition, data handling and analysis timescales. While we present data acquired with **ORIGAMI***^MS^* and then analysed by **ORIGAMI***^ANALYSE^*, the analysis module can be used as a standalone program on manually acquired datasets. We have shown applications most relevant to the field of structural biology, however the core functionality is not limited to large biomolecules; application for small molecule analysis where aIM-MS can be insightful is another area where this enhanced workflow may be useful.^39^ Since the software is open-source, written in Python and it utilises pre-existing Driftscope libraries, **ORIGAMI** is extensible we envisage additional functionality can be readily added to both **ORIGAMI***^MS^* and **ORIGAMI***^ANALYSE^*.

## Acknowledgements

L.G.M. would like to thank Emmy Hoyes for introduction to the wonderful world of WREnS scripting and Keith Richardson for initial help in the analysis of MassLynx raw files (Waters Corp., Manchester). We gratefully acknowledge BBSRC and MRC for support of this work in studentships to L.G.M. and A.F. respectively. This work was funded by the Biotechnology and Biological Sciences Research Council (BBSRC) by grants, BB/L015048/1 and BB/M017702/1. We also thank the British Mass Spectrometry Society (BMSS) and University of Manchester for continued support of our research.

## Author contributions

L.G.M. and B.B. designed **ORIGAMI***^MS^* whilst L.G.M. wrote both software components. L.G.M. and A.F. carried out the mass spectrometry experiments. L.G.M. analyse the data and wrote the manuscript. All authors contributed towards final editing of the manuscript.

## References

1. Lapthorn, C. et al. How useful is molecular modelling in combination with ion mobility mass spectrometry for ‘small molecule’ ion mobility collision cross-sections? Analyst (2015). doi:10.1039/C5AN00411J

2. Warnke, S. et al. Protomers of benzocaine: Solvent and permittivity dependence. J. Am. Chem. Soc. 137, 4236–4242 (2015).

3. Voronina, L. et al. Conformations of prolyl-peptide bonds in the bradykinin 1-5 fragment in solution and in the gas phase. (2016). doi:10.1021/jacs.6b04550

4. Beveridge, R. et al. Mass spectrometry locates local and allosteric conformational changes that occur on cofactor binding. Nat. Commun. 7, 12163 (2016).

5. Liu, Y., Cong, X., Liu, W. & Laganowsky, A. Characterization of Membrane Protein – Lipid Interactions by Mass Spectrometry Ion Mobility Mass Spectrometry. 579–586 (2017). doi:10.1007/s13361-016-1555-1

6. Clemmer, D. E., Hudgins, R. R. & Jamold, M. F. Naked Protein Conformations: Cytochrome. 10141–10142 (1995).

7. Quintyn, R. S., Zhou, M., Yan, J. & Wysocki, V. H. Surface-Induced Dissociation Mass Spectra as a Tool for Distinguishing Different Structural Forms of Gas-Phase Multimeric Protein Complexes. Anal. Chem. acs.analchem.5b03441 (2015). doi:10.1021/acs.analchem.5b03441

8. Pacholarz, K. J. et al. Hybrid Mass Spectrometry Approaches to Determine How L-Histidine Feedback Regulates the Enzyzme MtATP-Phosphoribosyltransferase. Structure 1–9 (2017). doi:10.1016/j.str.2017.03.005

9. Rabuck, J. N. et al. Activation state-selective kinase inhibitor assay based on ion mobility-mass spectrometry. Anal. Chem. 85, 6995–7002 (2013).

10. Harvey, S. R. et al. Small-molecule inhibition of c-MYC:MAX leucine zipper formation is revealed by ion mobility mass spectrometry. J. Am. Chem. Soc. 134, 19384–19392 (2012).

11. Pacholarz, K. J., Garlish, R. A., Taylor, R. J. & Barran, P. E. Mass spectrometry based tools to investigate protein-ligand interactions for drug discovery. Chem. Soc. Rev. 41, 4335–4355 (2012).

12. Jhingree, J. R. et al. Electron transfer with no dissociation ion mobility–mass spectrometry (ETnoD IM–MS). The effect of charge reduction on protein conformation. Int. J. Mass Spectrom. (2016). doi:10.1016/j.ijms.2016.08.006

13. Laszlo, K. J., Munger, E. B. & Bush, M. F. Folding of Protein Ions in the Gas Phase after Cation-to-Anion Proton-Transfer Reactions. J. Am. Chem. Soc. 138, 9581–9588 (2016).

14. Hopper, J. T. S. & Oldham, N. J. Collision induced unfolding of protein ions in the gas phase studied by ion mobility-mass spectrometry: the effect of ligand binding on conformational stability. J. Am. Soc. Mass Spectrom. 20, 1851–8 (2009).

15. Tian, Y. & Ruotolo, B. T. Ion Mobility-Mass Spectrometry and Collision Induced Unfolding Rapidly Detect Subtle Differences in Antibody Glycoforms Therapeutic Antibodies: Impact and Opportunity. (2016).

16. Mason, E. A. & McDaniel, E. W. Transport properties of ions in gases. (Wiley, New York, 1988).

17. Giles, K. et al. Applications of a travelling wave-based radio-frequency-only stacked ring ion guide. Rapid Commun. Mass Spectrom. 18, 2401–2414 (2004).

18. Giles, K., Williams, J. P. & Campuzano, I. Enhancements in travelling wave ion mobility resolution. Rapid Commun. Mass Spectrom. 25, 1559–1566 (2011).

19. Ruotolo, B. T., Benesch, J. L. P., Sandercock, A. M., Hyung, S.-J. & Robinson, C. V. Ion mobility-mass spectrometry analysis of large protein complexes. Nat. Protoc. 3, 1139–1152 (2008).

20. Niu, S. & Ruotolo, B. T. Collisional unfolding of multiprotein complexes reveals cooperative stabilization upon ligand binding. Protein Sci. 24, 1272–1281 (2015).

21. Eschweiler, J. D., Martini, R. M. & Ruotolo, B. T. Chemical Probes and Engineered Constructs Reveal a Detailed Unfolding Mechanism for a Solvent-Free Multidomain Protein. J. Am. Chem. Soc. jacs.6b11678 (2016). doi:10.1021/jacs.6b11678

22. Shelimov, K. B., Clemmer, D. E., Hudgins, R. R. & Jarrold, M. F. Protein Structure in Vacuo: Gas-Phase Conformations of BPTI and Cytochrome c. J. Am. Chem. Soc. 7863, 2240–2248 (1997).

23. Clemmer, D. E. & Jarrold, M. F. Ion Mobility Measurements and their Applications to Clusters and Biomolecules. J. Mass Spectrom. 32, 577–592 (1997).

24. Helden, G. Von, Gotts, N. & Bowers, M. T. Experimental evidence for the formation of fullerenes by collisional heating of carbon rings in the gas phase. Nature 363, 60–63 (1993).

25. Zhong, Y., Han, L. & Ruotolo, B. T. Collisional and Coulombic Unfolding of Gas-Phase Proteins: High Correlation to Their Domain Structures in Solution. Angew. Chem. Int. Ed. Engl. 53, 1–5 (2014).

26. Pagel, K., Hyung, S. J., Ruotolo, B. T. & Robinson, C. V. Alternate dissociation pathways identified in charge-reduced protein complex ions. Anal. Chem. 82, 5363–5372 (2010).

27. Tian, Y., Han, L., Buckner, A. C. & Ruotolo, B. T. Collision Induced Unfolding of Intact Antibodies: Rapid Characterization of Disulfide Bonding Patterns, Glycosylation, and Structures. Anal. Chem. acs.analchem.5b03291 (2015). doi:10.1021/acs.analchem.5b03291

28. Terral, G., Beck, A. & Cianf??rani, S. Insights from native mass spectrometry and ion mobility-mass spectrometry for antibody and antibody-based product characterization. J. Chromatogr. B Anal. Technol. Biomed. Life Sci. 1032, 79–90 (2016).

29. Zhao, Y. et al. Gas-Phase Analysis of the Complex of Fibroblast Growth Factor 1 with Heparan Sulfate: A Traveling Wave Ion Mobility Spectrometry (TWIMS) and Molecular Modeling Study. J. Am. Soc. Mass S 27, 1–14 (2016).

30. Eschweiler, J. D., Rabuck-Gibbons, J. N., Tian, Y. & Ruotolo, B. T. CIUSuite: A Quantitative Analysis Package for Collision Induced Unfolding Measurements of Gas-Phase Protein Ions. Anal. Chem. acs.analchem.5b03292 (2015). doi:10.1021/acs.analchem.5b03292

31. Allison, T. M. et al. Quantifying the stabilizing effects of protein-ligand interactions in the gas phase. Nat. Commun. 6, 8551 (2015).

32. Lima, I. Python for Scientific Computing Python Overview. Mar. Chem. 10–20 (2006). doi:10.1109/MCSE.2007.58

33. Hunter, J. D. Matplotlib: A 2D graphics environment. Comput. Sci. Eng. 9, 99–104 (2007).

34. Ramachandran, P. & Varoquaux, G. Mayavi: 3D visualization of scientific data. Comput. Sci. Eng. 13, 40–51 (2011).

35. Bush, M. F. et al. Collision cross sections of proteins and their complexes: a calibration framework and database for gas-phase structural biology. Anal. Chem. 82, 9557–9565 (2010).

36. Bush, M. F. Bush CCS Database. *http://depts.washington.edu/bushlab/ccsdatabase/* (2015).

37. Scarff, C. a, Thalassinos, K., Hilton, G. R. & Scrivens, J. H. Travelling wave ion mobility mass spectrometry studies of protein structure: biological significance and comparison with X-ray crystallography and nuclear magnetic resonance spectroscopy measurements. Rapid Commun. Mass Spectrom. 22, 3297–304 (2008).

38. Smith, D. P. et al. Deciphering drift time measurements from travelling wave ion mobility spectrometry-mass spectrometry studies. Eur. J. Mass Spectrom. (Chichester, Eng). 15, 113–130 (2009).

39. Gray, C. J. et al. Bottom-up elucidation of glycosidic bond stereochemistry. Anal. Chem. acs.analchem.6b04998 (2017). doi:10.1021/acs.analchem.6b04998

